# Spatiotemporal analysis reveals significant overlap of key proepicardial markers in the developing murine heart

**DOI:** 10.1101/666610

**Authors:** Irina-Elena Lupu, Andia N. Redpath, Nicola Smart

## Abstract

During embryonic development, the epicardium provides a source of multipotent progenitors for cardiac lineages, including pericytes, fibroblasts and coronary smooth muscle cells. The epicardium originates from a region of splanchnopleural mesoderm known as the proepicardial organ (PEO). The potential of the epicardium to contribute to coronary endothelium has been disputed, due to conflicting lineage tracing results with different PEO Cre lines. Controversy also surrounds when epicardial cell fate becomes restricted. Using single-cell RNA-sequencing, microscopy and flow cytometry-based single molecule RNA in situ hybridisation techniques, we systematically investigated the expression of five widely used epicardial markers, *Wt1, Tcf21, Tbx18, Sema3d* and *Scx*, over the course of development. We show co-expression of all markers in the PEO and epicardial layer until E13.5, then sequential downregulation as it undergoes quiescence. Markers also decrease in invading epicardium-derived progenitors, with the exception of *Tcf21*, lost only in epicardium-derived mural cells. Moreover, we demonstrate that the epicardium does not significantly contribute coronary endothelium. Our findings clarify a number of prevailing discrepancies in the field and support the notion that epicardial fate is not pre-determined within the PEO.

**Summary statement:** Assessing expression of five principal (pro)epicardial markers reveals their complete overlap during early embryonic development, challenging previous dogma regarding the existence of sub-compartments and the pre-committed fate model.

## Introduction

Extensive research has been dedicated to understanding how coronary endothelial cells (CECs) are derived during development. The established view of the PEO as the main origin of CECs has been challenged by the use of Cre-Lox genetic lineage tracing and by the recent demonstration that the main contributors of CECs in the developing heart are the sinus venosus and endocardium (Chen et al., 2014; Red-Horse et al., 2010; Tian et al., 2014; Tian et al., 2013; Zhang et al., 2016). However, a distinct proepicardial subcompartment was also proposed to give rise to CECs (Katz et al., 2012), raising questions about the vasculogenic potential of epicardial cells.

The PEO is a transient embryonic structure that, in mammals, arises in the septum transversum region (ST) from posterior second heart field progenitors (Kruithof et al., 2006; Lie-Venema et al., 2007). Proepicardial cells migrate onto the heart surface from embryonic day (E) 9.5 in mouse to form the outer epicardial layer. Epicardial cells undergo epithelial-to-mesenchymal transition (EMT) from E12.5, giving rise to a population of cells known as epicardium-derived cells (EPDCs). Whilst it is accepted that EPDCs further differentiate into pericytes (progenitors for coronary smooth muscle (Volz et al., 2015) and cardiac fibroblasts, their contribution to the coronary endothelium remains divisive, due to conflicting fate-mapping results (Acharya et al., 2012; Cai et al., 2008; Katz et al., 2012; Zhou and Pu, 2012). Cre-based lineage tracing, driven by the promoters of PEO genes *Wt1, Tbx18, Tcf21*, or by an enhancer of *Gata5*, reported little or no contribution to the coronary endothelium (Acharya et al., 2012; Cai et al., 2008; Merki et al., 2005; Zhou and Pu, 2012). In contrast, a distinct PEO subcompartment, expressing Sema3d and/or Scx, largely non-overlapping with previously identified markers, was reported to contribute: 7% of CECs at E16.5 from the *Sema3d* lineage and 25% of CECs were labelled postnatally with the *Scx* lineage (Katz et al., 2012). However, multiple studies report ubiquitous expression of *Wt1, Tcf21* and *Tbx18* in the early epicardium (Acharya et al., 2012; Wei et al., 2015), raising the question of whether non-overlapping subpopulations also exist at later stages in the epicardium proper. Another matter under scrutiny is whether EPDC fate specification is pre-determined within the PEO, or if these cells are multipotent.

In this report, we reveal the co-expression of all previously reported markers in the PEO and the entire epicardial layer early in development. We also provide evidence to suggest that epicardial cell fate is specified only after EMT and that EPDCs do not significantly contribute to coronary endothelial cells. Thus, our findings challenge previous conclusions, inferred by the use of non-specific Cre-based lineage tracing, around the existence of discrete epicardial sub-populations with pre-committed cell fates.

## Results & Discussion

### 1. PEO cells that transition to the epicardium co-express *Wt1, Sema3d, Tbx18, Scx* and *Tcf21*

We used single molecule RNA in situ hybridisation (RNAscope) on E9.5 sagittal mouse embryonic sections, to detect expression of *Wt1, Sema3d, Tcf21, Scx* and *Tbx18*, markers previously employed for PEO lineage tracing (Wang et al., 2012) (Fig. 1A-B). Three probes were multiplexed to allow simultaneous detection of the different markers. We observed complete overlap of marker expression in the PEO and in cells actively migrating towards the heart (Fig. 1C, Fig. S1A-B). Morphologically, proepicardial cells are defined as the protrusions/villi that extend from the ST region (Maya-Ramos et al., 2013). Based on our data, the *bona fide* PEO co-expresses all tested markers. Below the proepicardium, the markers labelled distinct domains of expression that only partially overlapped (Fig. S1A-B). Wt1 and Tcf21 are known to be expressed in the hepatic primordium, also located in the ST (Lu et al., 2000; Perez-Pomares et al., 2004), whereas Tbx18 expression is found in cardiomyocyte precursors present in the region (Christoffels et al., 2009).

**Fig.1.**
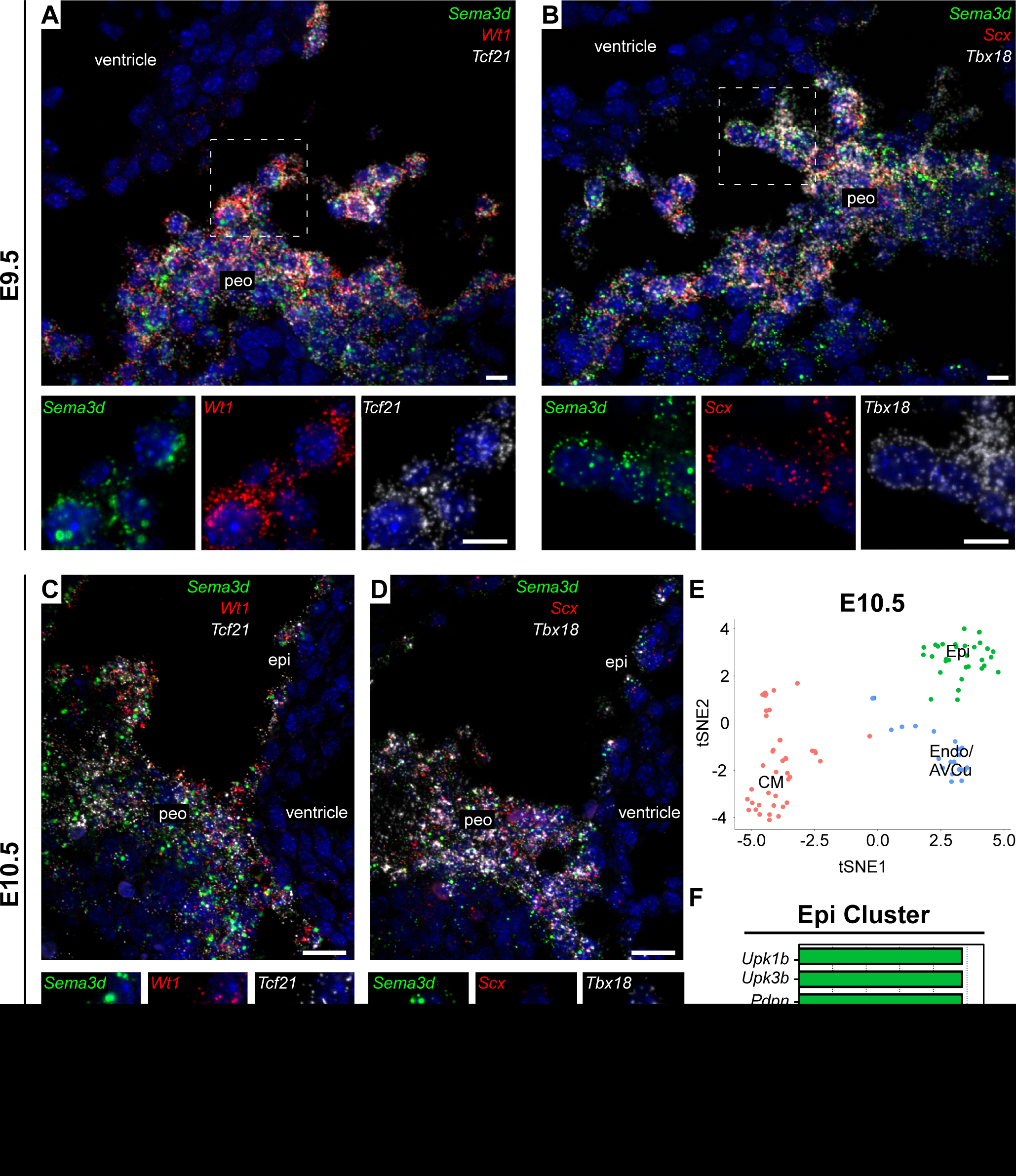
Complete overlap of (pro)epicardial markers *Sema3d, Wt1, Tcf21, Scx* and *Tbx18* in the proepicardium and founder epicardial cells. (A-B) RNA ISH staining of E9.5 embryo cryosections reveals co-expression of *Sema3d*, (A) *Wt1, Tcf21*, (B) *Scx*, and *Tbx18* mRNA in the proepicardial organ (n=5 embryos). (C-D) RNA ISH staining of E10.5 embryo cryosections demonstrate co-expression of *Sema3d*, (C) *Wt1, Tcf21*, (D) *Scx* and *Tbx18* mRNA in proepicardial cells migrating and subsequently forming the epicardium (n=2 embryos). (E) Two-dimensional tSNE plot demonstrating clustering of cardiomyocytes (CM), endocardium/atrioventricular cushion (Endo/AVCu), and epicardial cells (epi) from scRNA sequenced E10.5 hearts (total 96 cells; n=2 hearts).(F) Percentage of cells within the epicardial cluster (Epi) expressing selected epicardial genes. epi, epicardium; peo, proepicardial organ. Scale bars: 10µm.

To determine the marker profile of PEO cells that reach the heart and give rise to the definitive epicardium, we multiplexed RNAscope probes on E10.5 sagittal sections. Cells in contact with the heart surface co-expressed all markers (Fig. 1 C-D, Fig. S1A-B), although *Scx* expression decreased significantly compared to E9.5 (Fig. 1D). To independently confirm the RNAscope findings, we analysed published single-cell RNA-sequencing data from E10.5 mouse hearts (Dong et al., 2018). Clustering was performed using the significant principal components identified by Seurat, which classified the cells into three populations: epicardium (Epi), endocardium/atrioventricular cushion (Endo/AVCu) and cardiomyocytes (CM), shown here using the t-distributed stochastic neighbour embedding algorithm (tSNE) (Fig. 1E; Fig. S1C) (Butler et al., 2018; Platzer, 2013; Ratnadiwakara et al., 2018). *Wt1, Sema3d, Tcf21* and *Tbx18* were expressed in 97-100% of the epicardial cluster at E10.5, alongside mesothelial genes such as *Upk3b* and *Upk1b. Scx* was only expressed in 26% (Fig. 1F), but this may reflect the low capture rate of STRT-Seq technology (Ziegenhain et al., 2017).

Our findings show that proepicardial cells that transition onto the heart to form the epicardium co-express all the tested markers, contrasting with previous reports (Katz et al., 2012). The disparate findings most likely reflect a failure to distinguish the ST mesenchyme from the proepicardium. The ST mesenchyme is widely heterogeneous, containing precursors for a multitude of lineages, ranging from cardiomyocytes (Christoffels Vincent et al., 2006) to haematopoietic cells (Cañete et al., 2017). It is challenging to separate the proepicardium from the rest of the ST as there are no defined molecular boundaries and previously used markers, such as Wt1 or Tbx18, are also expressed in ST mesenchyme (Cai et al., 2008; Carmona et al., 2016). Our findings explain the differential potential of proepicardial/ST explants vs epicardial explants, with the former able to give rise to cardiomyocytes and endothelial cells, which are not present in epicardial outgrowths (Greulich and Kispert, 2013; Red-Horse et al., 2010; Ruiz-Villalba et al., 2013). We hypothesize that endothelial precursors, expressing markers such as Scx, might exist in the ST region and contribute to the heart, but transition via a non-epicardial route. Indeed, the proepicardial/ST markers used to infer coronary endothelial contribution, namely Scx, Sema3d and Gata4, also lineage-label cells in the SV/Endocardium (Cano et al., 2016; Katz et al., 2012), suggesting these precursors may contribute via the main CEC sources.

### 2. Wt1^CreERT2^ induction at E9.5 efficiently labels the epicardium

A key question is how the expression of epicardial markers changes throughout the course of development, both within the epicardial layer and in EPDCs as they differentiate. We used the inducible *Wt1CreERT2* (Zhou et al., 2008), crossed with *Rosa-tdTomato* reporter (Madisen et al., 2010), to label the early embryonic epicardium and enable tracing of the epicardial lineage. Maximal Cre recombination efficiency is reported to occur 24h-48h following tamoxifen delivery, therefore we administered tamoxifen at E9.5 in order to label the epicardium at E10.5-E11.5, before EMT and before *Wt1* is expressed in CECs, from E13.5 (Hayashi and McMahon, 2002; Zhou and Pu, 2012). RNAscope was performed at E11.5, multiplexing probes against epicardial markers and against *tdTomato* to determine extent of epicardial labelling (Fig. 2A-F). TdTomato efficiently labelled the epicardial layer, and careful inspection of high power images indicated an overlap with all epicardial markers throughout the entire epicardial layer (Fig. 2A-F). The only exception was a distinct population of *tdTomato*+ cells, located in the AV groove, which were positive for *Tcf21* (Fig. 2A, D). These cells derived from the *Wt1* lineage, as indicated by *tdTomato* expression, but down-regulated *Wt1, Tbx18, Sema3d* and *Scx* expression; Fig. 2A-F), suggesting the earliest transition to EPDCs from this region at E11.5. Some of the markers were detected in non-epicardial domains, *Tbx18* in cardiomyocytes (Fig. 2E) and *Scx* and *Tcf21* in AVCu (Fig. 2D, F), confirming previous reports (Acharya et al., 2011; Barnette et al., 2014; Christoffels et al., 2009).

**Fig.2.**
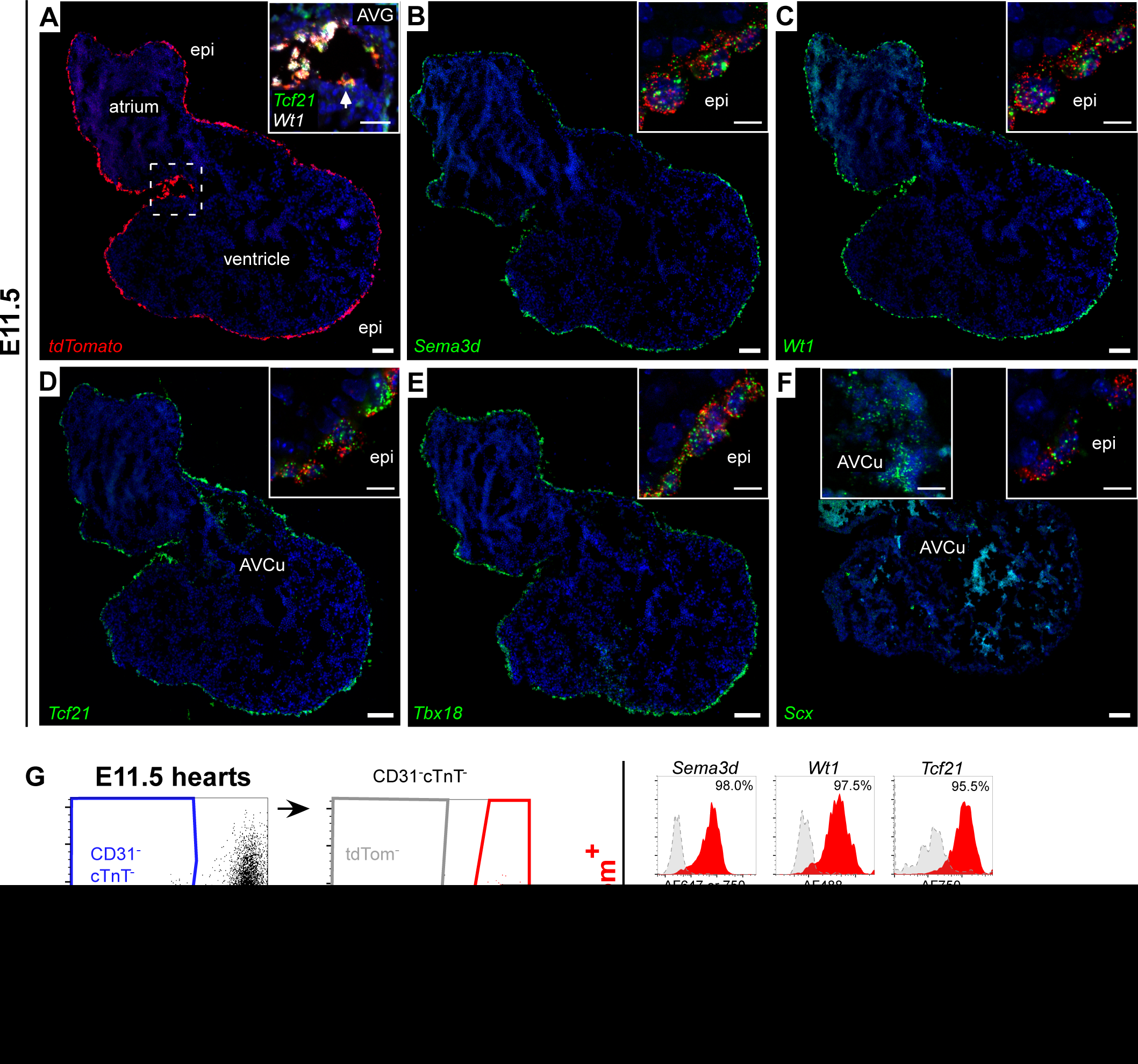
Substantial co-expression of epicardial markers in the epicardial layer. (A) RNA ISH staining of E11.5 heart cryosections shows tdTomato labelling of the epicardium, induced in Wt1^CreERT2^;Rosa26^tdTomato^embryos (n=3 hearts) at E9.5. Inset shows cells in atrioventricular groove (AVG) co-expressing *Tcf21* and *tdTomato*, which have downregulated *Wt1* (arrow). (B-F) RNA ISH staining of E11.5 heart cryosections reveals co-expression of (B) *Sema3d*, (C) *Wt1*, (D) *Tcf21*, (E) *Tbx18* and (F) *Scx* with *tdTomato*-labelled epicardium. (insets; n=3 hearts). (G) Flow cytometric analysis of dissociated E11.5 hearts. Epicardial cells, selected by gating CD31^-^cTnT^-^tdTomato^+^, show co-expression of *Sema3d, Wt1, Tcf21, Tbx18*, and *Scx* (n=31 hearts, representative of three independent experiments). Fluorescence minus one (FMO) control and percent positive shown. epi, epicardium; AVG, atrioventricular groove; AVCu, atrioventricular cushion. Scale bars: 100µm in A-F; 50µm in inset A, 10µm in inset B-F.

To quantitatively assess marker overlap at the single (whole) cell level, a flow cytometry-based RNA ISH method was utilised (PrimeFlow RNA Assay)(Lai et al., 2018) on enzymatically dissociated E11.5 hearts (Fig. 2G; Fig. S2D; with 90.6±0.2% WT1+ cells tdTomato labelled). After exclusion of cardiomyocytes and endothelial/endocardial cells, >95% of tdTomato+ cells co-expressed *Wt1, Tcf21, Sema3d* and *Tbx18. Scx* expression was very low and detected only in 88.7%. We concluded that inducing *Wt1CreERT2* at E9.5 efficiently labels the epicardium and its derivatives. We excluded the possibility that slight variations in *Wt1* expression might influence recombination efficiency by analysing non-lineage traced epicardial cells, which similarly co-expressed markers (e.g 97% and 92% Wt1+ cells expressed *Sema3d* and *Tcf21*, respectively; Fig. S1D). Expression levels measured for each marker fell within a limited dynamic range, as observed for the housekeeping gene *Actb* (Fig. S1D), and may reflect fluctuations associated with cell cycle or bursts in RNA production (Corrigan et al., 2016; Tirosh et al., 2016; Weinreb et al., 2018). Importantly, multiplexed probe sets were carefully selected to ensure highest assay channel sensitivity as fluorophore intensities between Alexa Fluor 488, 647 and 750 differ (Fig. S1D).

### 3. Epicardial markers are sequentially downregulated in the epicardium as development progresses

Next, we used the above labelling strategy to trace the epicardial lineage throughout the course of embryonic development, facilitated by *tdTomato* reporter expression (Fig. 3A). After tamoxifen delivery at E9.5, hearts were analysed at key developmental stages: E11.5, when epicardium formation is complete; E13.5, when epicardial cells invade the myocardium; E15.5, when epicardial EMT ceases and E17.5, when EPDCs have differentiated. The expression of each epicardial marker against *tdTomato* reporting was assessed at every stage by RNAscope and Primeflow (Fig. 3, Fig. S2A-E, Fig. S3A). By later stages, we detected widespread expression of epicardial markers in other cell types, highlighting the limitations of constitutive Cre lines (Fig. S2A-E). *Sema3d* and *Scx* were expressed in the AVCu (Fig. S2A, D) (Katz et al., 2012); *Tbx18* expression was found in vascular smooth muscle cells (vSMCs), both of non-epicardial (tdTomato-aortic vSMCs; Fig. S3B) and epicardial origin (tdTomato+ coronary vSMCs; Fig. S3C), from E15.5, as well as throughout the left ventricle (Fig. S2E; Fig. S3D); *Wt1* was expressed in the endothelium after E13.5 (Fig. S2B; Fig. S3E-G) and *Tcf21* in the interstitial cells (Fig. S2C), confirming other published data (Acharya et al., 2011; Zhou and Pu, 2012). Due to the non-specificity of these markers, we focused on tdTomato+ cells, representing the epicardial lineage, for our subsequent analyses.

**Fig.3.**
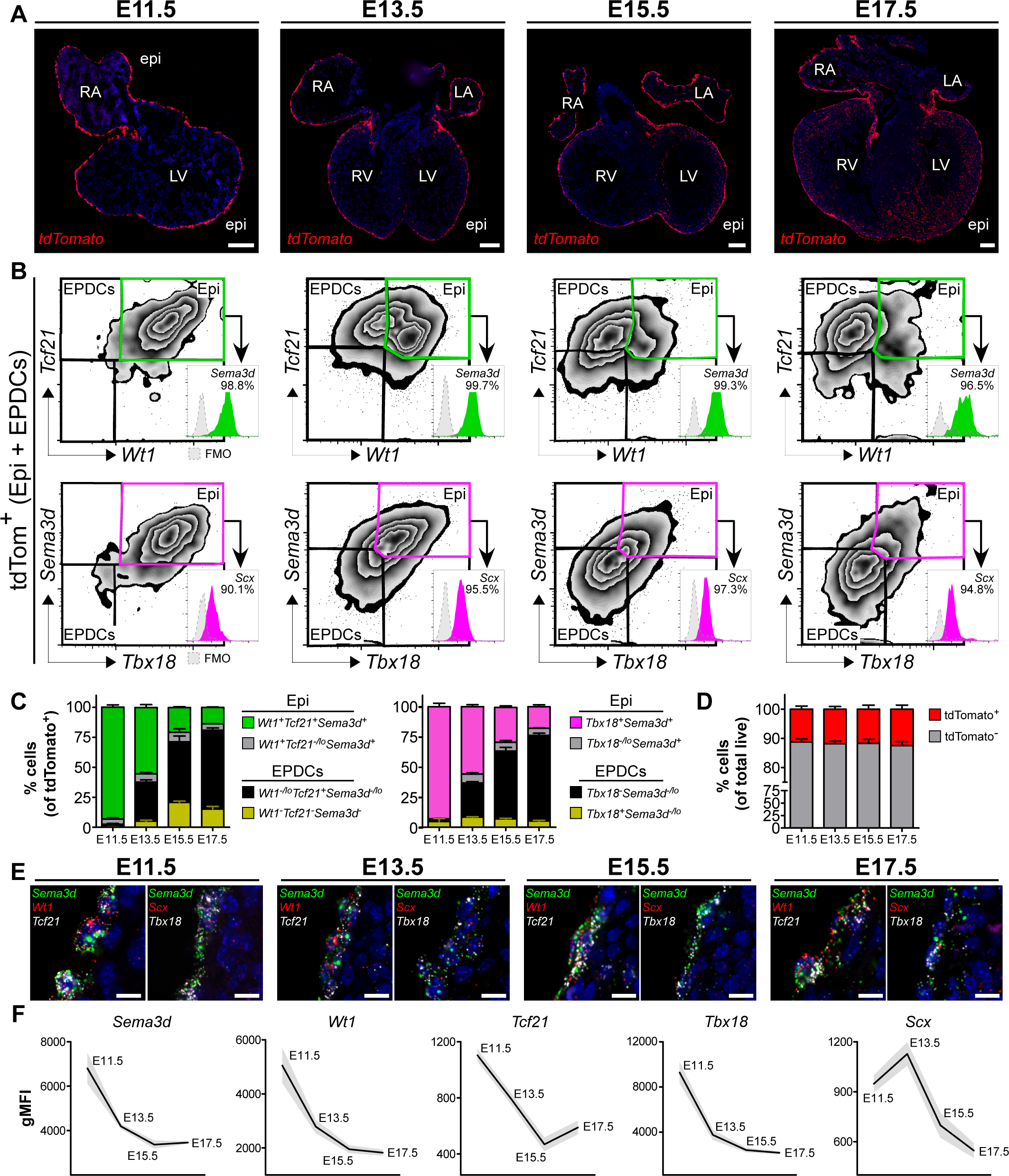
Epicardial markers are downregulated in invading epicardium-derived cells whilst the epicardium largely retains its signature. (A) RNA ISH staining of E11.5, E13.5, E15.5 and E17.5 heart cryosections shows tdTomato labelling of the epicardium and lineage-traced EPDCs (n=3 hearts/stage). (B) Flow cytometric analysis of dissociated E11.5 (n=31 hearts), E13.5 (n=25 hearts; representative of three independent experiments, E15.5 (n=10 hearts) and E17.5 hearts (n=2 hearts; representative of two independent experiments).The epicardial lineage, selected by gating CD31^-^cTnT^-^tdTomato^+^, shows overall loss of *Sema3d, Wt1, Tcf21, Tbx18*, and *Scx* as development progresses due to the increasing production of EPDCs. TdTomato^+^ cells co-expressing *Sema3d, Wt1, Tcf21, Tbx18*, and *Scx* (green and purple) represent the epicardium. Fluorescence minus one (FMO) control and percent positive shown. (C) Populations expressing *Sema3d, Wt1, Tcf21, Tbx18*, and *Scx*, calculated as a proportion of tdTomato^+^ cells, demonstrate gradual expansion of the EPDC fraction as development progresses. (n= as Fig3B; data shown as mean ± s.e.m of at least two independent experiments). (D) Flow cytometric analysis of the proportion of tdTomato^+^ cells during development (E11.5 n=31; E13.5 n=25; E15.5 n=19; E17.5 n=9; data shown as mean ± s.e.m of three independent experiments). (E) RNA ISH staining of E11.5, E13.5, E15.5 and E17.5 heart cryosections shows co-expression of *Sema3d, Wt1, Tcf21, Tbx18*, and *Scx* in cells retained in the epicardium. (F) Flow cytometric analysis of *Sema3d, Wt1, Tcf21, Tbx18, and Scx* in the epicardial population demonstrates a rapid decrease in their expression levels between E11.5 and E15.5 (data shown as mean gMFI ± s.e.m of four independent experiments with exception of Tbx18 N=2). gMFI, geometric Mean Fluorescence Intensity. RA, right atrium; RV, right ventricle; LA, left atrium; LV, left ventricle; Epi, epicardium; AVCu, atrioventricular cushion. Scale bars: 200µm in A; 10µm in E.

To assess co-expression of markers in the epicardial lineage across these stages, two different probe combinations were used for PrimeFlow on tdTomato+ hearts: *Sema3d/Wt1/Tcf21* and *Sema3d/Scx/Tbx18* with *Sema3d* serving to ensure parity across probe sets(Fig. 3B, gating only on tdTomato+ cells, Fig. S3A), and the proportion of tdTomato+ cells expressing each marker was quantified at every stage (Fig. 3C). At E11.5, tdTomato+ cells almost exclusively (>95%) represented the epicardium, which continued to co-express all marker genes (Fig. 3A-C, Fig. S3A). From E13.5, we detected the emergence of EPDCs, as shown by tdTomato+ cells present within the myocardium, and by the appearance of a distinct additional population of cells in the flow cytometry scatter plots, which had decreased expression of *Sema3d, Wt1, Tbx18 and Scx* (Fig. 3B, Fig. S3A). The number of EPDCs increased as development progressed, exceeding the number of cells within the epicardial layer (Fig.3A-C), however, the combined number of epicardial and epicardium-derived cells, as a proportion of the total heart cells, remained constant throughout development, at around 12% (Fig. 3D). The expression of all epicardial markers was downregulated, and eventually lost, in EPDCs, with the exception of *Tcf21*, which increased (Fig. 3B). All EPDCs were positive for *Tcf21* at E13.5, implying that *Tcf21* expression is not sufficient to determine cell fate. From E15.5, Tcf21 was downregulated in a subpopulation of EPDCs, coinciding with the appearance of epicardium-derived mural cells (Volz et al., 2015), (Fig. 3B).

Expression of *Sema3d, Wt1, Tcf21* and *Tbx18* peaked within the epicardium at E11.5, as assessed by both RNAscope and PrimeFlow (Fig. 3E-F). *Tcf21* expression was dramatically reduced beyond E13.5 in the epicardium, being almost undetectable in some of the cells, followed by a reduction in *Tbx18* expression (Fig. 3E, Fig. S4A-B), to coincide with completion of epicardial EMT by E15.5 (Liu et al., 2016), and onset of epicardial quiescence. *Wt1* and *Sema3d* continued to be co-expressed in the entire epicardium throughout embryonic development, albeit at lower levels (Fig. 3F; Fig. S4A-B). Although markers were sequentially downregulated, marker expression remained discernibly homogeneous between individual epicardial cells; thus, our study found no evidence of discrete sub-populations, at least at the level of the markers tested.

### 4. The epicardium produces multipotent progenitors that contribute to cardiac fibroblasts and mural cells, but not to endothelium

To further investigate the epicardial lineage in greater detail, we performed single-cell RNA-sequencing of tdTomato+ FACS sorted cells at E15.5, a stage when derivative fates manifest (Picelli et al., 2014). Five transcriptionally-distinct populations were identified using the significant principal components and visualized with tSNE, as facilitated by Seurat (Butler et al., 2018; Stuart et al., 2018) (Fig. 4A). The first cluster represented the epicardium (Epi), based on expression of mesothelial genes, such as Upk3b (Fig. S4C). Two mesenchymal clusters were identified, the first (Mes1) presented a transcriptional profile consistent with the subepicardial mesenchyme reported by Xiao et al., whereas the second cluster (Mes2) was more mature, expressing genes such as *Postn* (Xiao et al., 2018). A proliferating cluster was also identified (Pro), bearing a transcriptional signature similar to the Mes2 cluster. The fifth cluster represented the epicardium-derived mural cells and expressed multiple pericyte-associated genes, such as *Cspg4* or *Rgs5*. We used a dot plot to analyse expression of epicardial markers within the different clusters (Fig. 4B-C). *Sema3d* and *Wt1* were expressed in the entire epicardial cluster (100%; Fig.4B), whereas their expression was lower in the mesenchymal populations and almost undetectable in the mural cluster. Levels of *Tcf21* in the epicardium were very low, however expression was high in both mesenchymal clusters. *Tbx18* was mainly localised to the epicardium and the mural cell cluster, whereas *Scx* was detected in 69% of the epicardial cluster (Fig. 4B-C), likely due to its very low expression.

**Fig.4.**
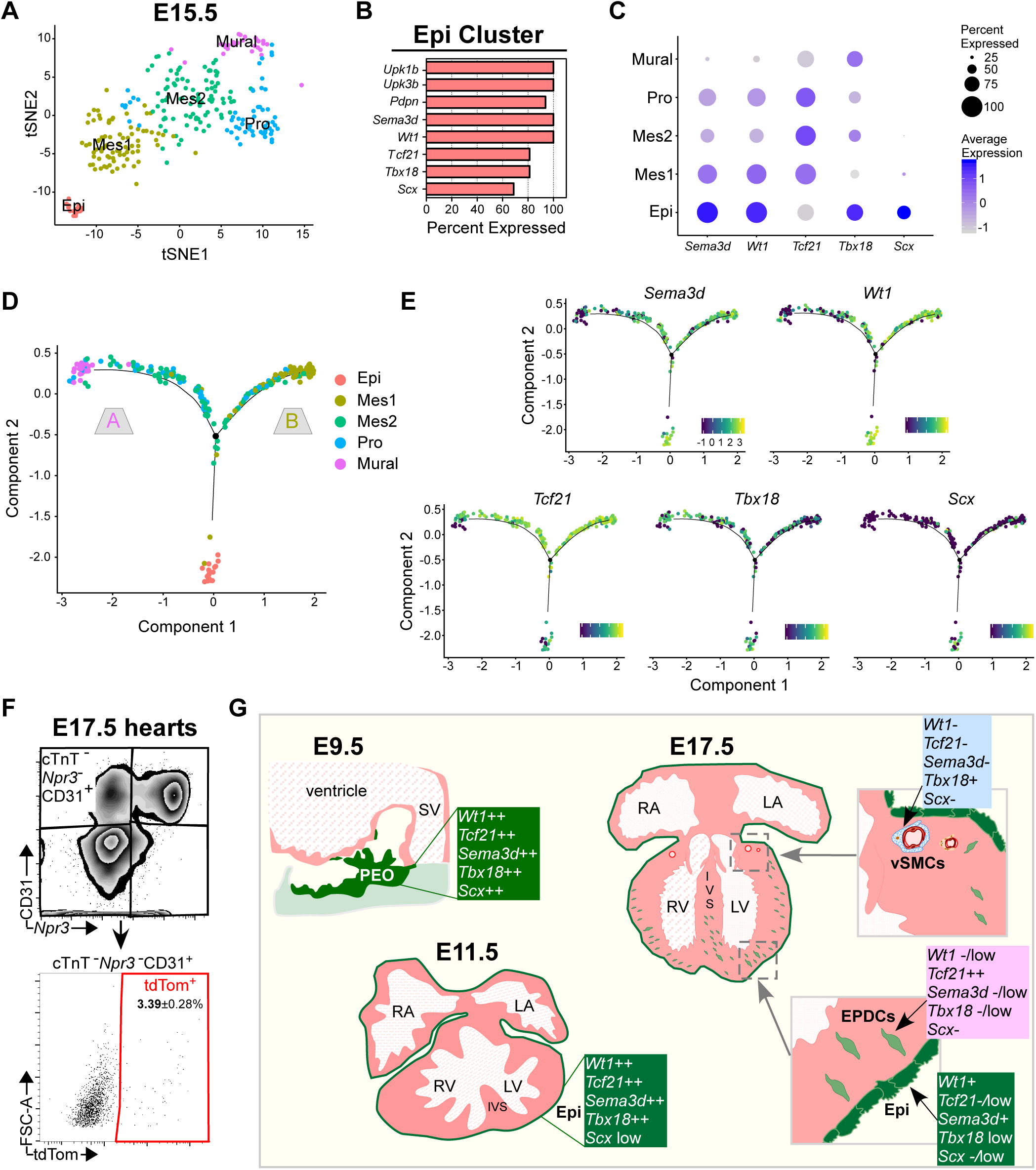
scRNAseq profiling demonstrates the epicardium is a source of fibroblasts and mural cells, and cell fate is not pre-determined by a single epicardial marker. (A) Two-dimensional tSNE plot demonstrating clustering of epicardial (Epi), mesenchymal (Mes1 and Mes2), mural and proliferating (Pro) cells derived from scRNA sequenced tdTomato^+^ FACS sorted cells at E15.5 (total 276 cells; n=6 hearts). (B) Percentage of cells within the epicardial cluster (Epi) expressing selected epicardial genes. (C) Dot plot showing proportion of cells in each cluster expressing selected epicardial genes. Dot size represents the percentage of cells expressing, and the colour scale indicates the average expression level. (D) Pseudotime trajectory of tdTomato^+^ cells at E15.5, showing bifurcation to either branch A (mural) or branch B (mesenchymal 1). Cells are coloured by cluster identity. (E) Pseudotime trajectory coloured by expression level of selected epicardial genes. (F) Flow cytometric analysis of dissociated E17.5 hearts (Wt1^CreERT2^; Rosa^26tdTomato^). Endothelial cells were selected by gating cTnT^-^Npr3^-^CD31^+^ and downstream tdTomato^+^ gating to determine epicardial contribution. Percentage of tdTomato^+^ endothelial cells expressed as mean ± s.e.m of six biological replicates (n=6 hearts). (G) Schematic summarising marker profile of epicardium and its derivatives across the time course of development.

We used Monocle2 to classify cells based on pseudotime, in order to reconstruct the differentiation trajectory of the epicardial lineage (Qiu et al., 2017a; Qiu et al., 2017b; Trapnell et al., 2014). The developmental trajectory obtained was very similar to that reported by Xiao et al., with one branching point leading towards differentiated cells (branch A) and one consisting of more mesenchymal cells (branch B) (Fig. 4D) (Xiao et al., 2018). In our case, branch A was associated with expression of mural genes, such as *Pdgfrβ* and *Rgs5*, whereas branch B had higher expression of fibroblast-like genes, such as *Pdgfra* and *Dpt* (Fig. S4D). Expression of *Sema3d, Wt1, Tcf21, Tbx18 and Scx* was visualised across the trajectory. None of the genes showed branch-dependent expression, suggesting their expression is not sufficient to restrict cell fate (Fig. 4E). The branch point occurred after mesenchymal cells first emerged, suggesting that cell fate may be specified in EPDCs post-EMT. This model contradicts a previous report, which suggested that fate restriction of EPDCs occurred prior to EMT (Acharya et al., 2012). Tcf21 was concluded to be required only for EMT of epicardial cells destined to become fibroblasts, because PDGFRa+, but not PDGFRβ+, cells were absent from Tcf21 global knock out embryos. However, another study showed that Tcf21 is required for normal formation of the entire epicardium, raising questions about the ability of Tcf21-depleted epicardial cells to undergo EMT altogether (Tandon et al., 2013). Recently, the endocardium was identified as an alternative source of mural cells in the heart, therefore it is possible the PDGFRβ+ cells present in Tcf21 mutants derived from this compartment (Chen et al., 2016). This controversy could be resolved in future by conditional deletion of Tcf21 and lineage tracing with Wt1^CreERT2^ to assess the origin of the mural cells in these hearts.

In addition to the single-cell RNA-sequencing, we also performed flow cytometric analysis at E17.5 to quantify epicardial contribution to the coronary endothelium, labelled with CD31 (excluding endocardial cells by *Npr3* expression (Zhang et al., 2016)). Of the CD31+*Npr3*-population, tdTomato+ cells averaged just 3.4% (Fig. 4F). Wt1-lineage contribution to CECs seems to be negligible, which is in agreement with previous reports (Cai et al., 2008; Zhou and Pu, 2012).

In conclusion, our data show an overlap in expression of the epicardial markers *Wt1, Tcf21, Sema3d, Tbx18* and *Scx*, in the epicardium early in development, with minimal variation in level of expression between cells. We postulate that previous, contrasting (non-inducible) lineage tracing results may reflect expression of the Cre-driver in non-epicardial lineages, rather than the existence of subpopulations within the epicardial layer. Genes outside the tested ‘canonical epicardial signature’ may be heterogeneously expressed (Bollini et al., 2014; Cao et al., 2016), however, based on single-cell RNA-sequencing inferred developmental trajectories, we suggest that the fate of epicardium-derived cells is specified after EMT, potentially due to extrinsic signalling (Schematic, Fig. 4G).

## Materials & Methods

### Mouse strains

*Wt1*^*CreERT2*^ (Zhou et al., 2008), *Rosa26*^*tdTomato*^(Madisen et al., 2010) alleles have been previously described. Males homozygous for the *Rosa26*^*tdTomato*^ allele and heterozygous for *Wt1*^*CreERT2*^ were crossed with C57BL/6 females to generate embryos. The mice used were on a C57BL/6 genetic background. Pregnant females were oral gavaged with 80mg/kg tamoxifen (Sigma, #T5648) 9 days after the mating plug was detected. Tamoxifen was dissolved in peanut oil with 10% ethanol at a final concentration of 10mg/ml. The animals were kept in a controlled environment and all procedures used were approved by the University of Oxford Animal Welfare and Ethical Review Boards in accordance with Animals (Scientific Procedures) Act 1986 (Home Office, United Kingdom).

### RNAscope & Immunofluorescence

RNAscope is a commercially available in-situ hybridization assay provided by Advanced Cell Diagnostics (ACD). RNAscope was performed on 8-12 µm thick embryonic cryosections that had been fixed for 2h in 4%PFA and embedded in OCT (Tissue-Tek). The following catalogue probes were used: Wt1 #432711), Sema3d (#488111), Tcf21 (#508661), Tbx18 (#515221), Scx (#439981), tdTomato (#317041). The 3-plex negative probe against *dapB* was used as a negative control (#320878). The RNAscope Multiplex Fluorescent Reagent Kit v2 assay was performed according to manufacturer’s instructions, except for the target retrieval step which was reduced to 10min and the Protease Plus digestion which was performed at room temperature (RT). The TSA plus fluorophores (PerkinElmer, NEL753001KT, NEL745E001KT) were used at 1:500 final concentration for Fluorescein, 1:1000 for Cy3 and 1:1500 for Cy5.

For immunostaining the cryosections were rehydrated for 15 min in PBS, then permeabilised in PBS + 0.5%triton for 15 min at RT, followed by blocking in PBS + 10% goat serum +1% bovine serum albumin (BSA) for 1h at RT and incubation with primary antibody diluted in blocking buffer overnight at 4°C. The sections were rinsed three times, then washed three times in PBS + 0.1% Triton X-100, then the secondary antibodies diluted in blocking buffer were added for 1 hour, RT. The sections were rinsed three times in PBS containing 0.1% Triton X-100, then washed twice for 5 min/each wash, followed by 5 min incubation with DAPI nuclear stain. The slides were mounted in Vectashield (Vector Laboratories, H-1000). Antibodies used: WT1 1:100 dilution (Abcam, ab89901), CD31/Pecam1 1:100 dilution (Abcam, ab119341).

Sections were imaged using a Leica DM6000 fluorescence microscope and Olympus Fluoview-1000 confocal microscope and processed using Fiji.

### PrimeFlow RNA Assay & Flow Cytometry

Hearts were enzymatically dissociated using the Neonatal Heart Dissociation Kit (#130-098-373) and the gentleMACS Octo Dissociator with heaters (#130-096-427) from Milteny Biotec according to manufacturer’s instructions. The dissociated hearts were further processed using the PrimeFlow RNA Assay Kit from ThermoFisher Scientific (#88-18005). Briefly, cells were surface stained with the Zombie Aqua Fixable Viability Kit (Biolegend, #423101) for 0.5hr at RT,1:1000 dilution, followed by BV421 rat anti-CD31 antibody (Biolegend, #102423) for 0.5hr on ice, 1:50 dilution. Samples were further fixed and permeabilised using reagents provided in the kit. Intracellular staining was performed using BV421 mouse anti-cardiac troponin T antibody (BD Bioscieces, #565618) for 0.5hr on ice, 1:200 dilution, followed by further fixation and hybridization as specified in the instructions. PrimeFlow catalogue probes were used against Scx (#VB1-3028071-PF), Tbx18 (#VB4-3113124-PF), Tcf21 (#VB6-3197774-PF), Sema3d (#VB1-3044712-PF), Wt1 (#VB4-13886-PF) and Npr3 (VB1-3031000-PF). Probes against dapb (#VF1-11712-PF, # VF6-10407-PF, # VF4-10408-PF) were used as negative controls and probes against Actb (#VB6-12823-PF, # VB1-10350-PF, # VB4-10432-PF) as positive controls.

For analysing the contribution of the epicardial lineage to endothelium hearts were dissociated and surface stained with viability dye as described above, then pre-treated for 5 min on ice with TruStain FcX (anti-mouse CD16/32) antibody (Biolegend, #101319), 1:50 to block Fcγ receptors. Cells were subsequently stained with BV605 rat anti-CD31 antibody, (Biolegend, #102427) on ice for 0.5hr, 1:50 dilution. Cells were fixed and permeabilised using the PrimeFlow kit reagents, then intracellularly stained with cTNT (as described above). Cells were stored in IC fixation buffer (ThermoFisher, #00-8222-49) until acquisition.

All samples were analysed using BD LSRFORTESSA X-20 cytometer and processed using FlowJo software. Percent positive was calculated using Super-Enhanced Dmax Subtraction algorithm (%SED;FlowJo) when comparing histograms (Supplementary information Table S1-S3). Geometric mean fluorescence intensity (gMFI) was calculated by subtracting fluorescence minus one (FMO) gMFI from sample gMFI.

### Single-cell RNA-sequencing & Analysis

E10.5 single-cell heart transcriptome data was downloaded as TPM from Gene Expression Omnibus (GSM3027035) (Dong et al., 2018). The TPM values were log10 transformed and analysed using Seurat in R (http://www.R-project.org/ (Stuart et al., 2018). In brief, the significant principal components were used to classify the cells into clusters and the tSNE method was used to visualise the clusters (Platzer, 2013).

For the E15.5 single-cell RNA-Sequencing data, the ventricles from 6 embryonic hearts were dissociated (as described above) and BD FACSAria III was used to sort cells positive for tdTomato fluorescence into 96-well plates. The cells were processed according to the Smart-Seq2 protocol (Picelli et al., 2014). Tagmentation and library preparation was done using the Nextera XT DNA Library Prep kit and sequenced using the NextSeq 500/550 High Output Kit v2 (75 cycles) 400 million reads (Illumina, #FC-404-2005) on Illumina NextSeq 500 platform. BCL files from the sequencer were converted to FastQ with bcl2fastq (Illumina, v.2.19.1.403) using default settings. Samples with fewer reads in total than the empty wells were discarded. Reads were trimmed using default settings and “--nextera” for adapter clipping (TrimGalore v.0.4.4). Random samples were checked with FastQC (v.0.11.3) to control for the quality of the sequence. Reads were aligned to the mouse genome (GRCm38/Mm10) with STAR (v.2.3.3a) in gene counting mode with Gencode (v.M16) annotations minus the megatranscript Gm20388 (Dobin et al., 2013). Splice junctions from all samples were combined after the 1st pass alignment. Non-canonical junctions and junctions covered with < 10 reads were removed prior to running the 2nd pass alignment. The samples were further processed with Seurat. Genes with > 200 reads in at least 3 cells were retained for downstream processing. Cells were filtered to retain those with < 5% mitochondrial genes and at least 500 expressed genes. 276 cells passed this threshold. Reads were normalized to sequencing depth and scaled. Cell cycle scores (G2M and S) was predicted using G2M and S phase genes (Kowalczyk et al., 2015). The difference between the G2M and S phase score was regressed out using the cell cycle regression Seurat vignette. Clustering was performed as described above. The normalised data from Seurat was imported into Monocle2 for analysis of pseudotime (Qiu et al., 2017a) (Qiu et al., 2017b).

## Acknowledgements

We thank Professor William Pu, Harvard Medical School, for the *Wt1*CreERT2 line; Dr Madeleine Lemieux (Bioinfo) for processing the SMART-Seq2 data; the WIMM Single cell genomics facility (University of Oxford), especially Dr Neil Ashley, for producing the SMART-Seq2 libraries and sequencing the samples, the Sir William Dunn School of Pathology Flow Cytometry facility for providing helpful advice, Micron for microscopy facilities and the Biomedical Services Staff for animal husbandry.

## Author Contributions

Conceptualisation: I-EL, NS; Experimental design: I-EL, ANR, NS; Data acquisition and analysis: I-EL, ANR; Manuscript (initial draft): I-EL; Manuscript (figure preparation): ANR; Manuscript (editing): ANR, NS; Bioinformatic analysis: IE-L, ANR; Supervision: ANR, NS; Funding acquisition: NS.

## References

Acharya, A., Baek, S. T., Banfi, S., Eskiocak, B. and Tallquist, M. D. (2011). Efficient inducible Cre-mediated recombination in Tcf21cell lineages in the heart and kidney. genesis 49, 870–877.

Acharya, A., Baek, S. T., Huang, G., Eskiocak, B., Goetsch, S., Sung, C. Y., Banfi, S., Sauer, M. F., Olsen, G. S., Duffield, J. S., et al. (2012). The bHLH transcription factor Tcf21 is required for lineage-specific EMT of cardiac fibroblast progenitors. Development 139, 2139–2149.

Barnette, D. N., VandeKopple, M., Wu, Y., Willoughby, D. A. and Lincoln, J. (2014). RNA-Seq Analysis to Identify Novel Roles of Scleraxis during Embryonic Mouse Heart Valve Remodeling. PLOS ONE 9, e101425.

Bollini, S., Vieira, J. M., Howard, S., Dube, K. N., Balmer, G. M., Smart, N. and Riley, P. R. (2014). Reactivated adult epicardial progenitor cells are a heterogeneous population molecularly distinct from their embryonic counterparts. Stem Cells Dev 23, 1719–1730.

Butler, A., Hoffman, P., Smibert, P., Papalexi, E. and Satija, R. (2018). Integrating single-cell transcriptomic data across different conditions, technologies, and species. Nat Biotechnol 36, 411–420.

Cai, C. L., Martin, J. C., Sun, Y., Cui, L., Wang, L., Ouyang, K., Yang, L., Bu, L., Liang, X., Zhang, X., et al. (2008). A myocardial lineage derives from Tbx18 epicardial cells. Nature 454, 104–108.

Cañete, A., Carmona, R., Ariza, L., Sánchez, M. J., Rojas, A. and Muñoz-Chápuli, R. (2017). A population of hematopoietic stem cells derives from GATA4-expressing progenitors located in the placenta and lateral mesoderm of mice. Haematologica 102, 647.

Cano, E., Carmona, R., Ruiz-Villalba, A., Rojas, A., Chau, Y.-Y., Wagner, K. D., Wagner, N., Hastie, N. D., Muñoz-Chápuli, R. and Pérez-Pomares, J. M. (2016). Extracardiac septum transversum/proepicardial endothelial cells pattern embryonic coronary arterio–venous connections. Proceedings of the National Academy of Sciences of the United States of America 113, 656–661.

Cao, J., Navis, A., Cox, B. D., Dickson, A. L., Gemberling, M., Karra, R., Bagnat, M. and Poss, K. D. (2016). Single epicardial cell transcriptome sequencing identifies Caveolin 1 as an essential factor in zebrafish heart regeneration. Development 143, 232–243.

Carmona, R., Canete, A., Cano, E., Ariza, L., Rojas, A. and Munoz-Chapuli, R. (2016). Conditional deletion of WT1 in the septum transversum mesenchyme causes congenital diaphragmatic hernia in mice. Elife 5.

Chen, H. I., Sharma, B., Akerberg, B. N., Numi, H. J., Kivelä, R., Saharinen, P., Aghajanian, H., McKay, A. S., Bogard, P. E., Chang, A. H., et al. (2014). The sinus venosus contributes to coronary vasculature through VEGFC-stimulated angiogenesis. Development (Cambridge, England) 141, 4500–4512.

Chen, Q., Zhang, H., Liu, Y., Adams, S., Eilken, H., Stehling, M., Corada, M., Dejana, E., Zhou, B. and Adams, R. H. (2016). Endothelial cells are progenitors of cardiac pericytes and vascular smooth muscle cells. Nature Communications 7, 12422.

Christoffels Vincent, M., Mommersteeg Mathilda, T. M., Trowe, M.-O., Prall Owen, W. J., de Gier-de Vries, C., Soufan Alexandre, T., Bussen, M., Schuster-Gossler, K., Harvey Richard, P., Moorman Antoon, F. M., et al. (2006). Formation of the Venous Pole of the Heart From an Nkx2–5–Negative Precursor Population Requires Tbx18. Circulation Research 98, 1555–1563.

Christoffels, V. M., Grieskamp, T., Norden, J., Mommersteeg, M. T., Rudat, C. and Kispert, A. (2009). Tbx18 and the fate of epicardial progenitors. Nature 458, E8–9; discussion E9-10.

Corrigan, A. M., Tunnacliffe, E., Cannon, D. and Chubb, J. R. (2016). A continuum model of transcriptional bursting. Elife 5.

Dobin, A., Davis, C. A., Schlesinger, F., Drenkow, J., Zaleski, C., Jha, S., Batut, P., Chaisson, M. and Gingeras, T. R. (2013). STAR: ultrafast universal RNA-seq aligner. Bioinformatics (Oxford, England) 29, 15–21.

Dong, J., Hu, Y., Fan, X., Wu, X., Mao, Y., Hu, B., Guo, H., Wen, L. and Tang, F. (2018). Single-cell RNA-seq analysis unveils a prevalent epithelial/mesenchymal hybrid state during mouse organogenesis. Genome Biol 19, 31.

Greulich, F. and Kispert, A. (2013). Epicardial Lineages. Journal of Developmental Biology 1, 32.

Hayashi, S. and McMahon, A. P. (2002). Efficient recombination in diverse tissues by a tamoxifen-inducible form of Cre: a tool for temporally regulated gene activation/inactivation in the mouse. Dev Biol 244, 305–318.

Katz, T. C., Singh, M. K., Degenhardt, K., Rivera-Feliciano, J., Johnson, R. L., Epstein, J. A. and Tabin, C. J. (2012). Distinct Compartments of the Proepicardial Organ Give Rise to Coronary Vascular Endothelial Cells. Developmental Cell 22, 639–650.

Kowalczyk, M. S., Tirosh, I., Heckl, D., Rao, T. N., Dixit, A., Haas, B. J., Schneider, R. K., Wagers, A. J., Ebert, B. L. and Regev, A. (2015). Single-cell RNA-seq reveals changes in cell cycle and differentiation programs upon aging of hematopoietic stem cells. Genome research 25, 1860–1872.

Kruithof, B. P., van Wijk, B., Somi, S., Kruithof-de Julio, M., Perez Pomares, J. M., Weesie, F., Wessels, A., Moorman, A. F. and van den Hoff, M. J. (2006). BMP and FGF regulate the differentiation of multipotential pericardial mesoderm into the myocardial or epicardial lineage. Dev Biol 295, 507–522.

Lai, C., Stepniak, D., Sias, L. and Funatake, C. (2018). A sensitive flow cytometric method for multiparametric analysis of microRNA, messenger RNA and protein in single cells. Methods 134-135, 136–148.

Lie-Venema, H., van den Akker, N. M., Bax, N. A., Winter, E. M., Maas, S., Kekarainen, T., Hoeben, R. C., deRuiter, M. C., Poelmann, R. E. and Gittenberger-de Groot, A. C. (2007). Origin, fate, and function of epicardium-derived cells (EPDCs) in normal and abnormal cardiac development. Scientific World Journal 7, 1777–1798.

Liu, Q., Zhang, H., Tian, X., He, L., Huang, X., Tan, Z., Yan, Y., Evans, S. M., Wythe, J. D. and Zhou, B. (2016). Smooth muscle origin of postnatal 2nd CVP is pre-determined in early embryo. Biochemical and biophysical research communications 471, 430–436.

Lu, J., Chang, P., Richardson, J. A., Gan, L., Weiler, H. and Olson, E. N. (2000). The basic helix-loop-helix transcription factor capsulin controls spleen organogenesis. Proc Natl Acad Sci U S A 97, 9525–9530.

Madisen, L., Zwingman, T. A., Sunkin, S. M., Oh, S. W., Zariwala, H. A., Gu, H., Ng, L. L., Palmiter, R. D., Hawrylycz, M. J., Jones, A. R., et al. (2010). A robust and high-throughput Cre reporting and characterization system for the whole mouse brain. Nature neuroscience 13, 133–140.

Maya-Ramos, L., Cleland, J., Bressan, M. and Mikawa, T. (2013). Induction of the Proepicardium. J Dev Biol 1, 82–91.

Merki, E., Zamora, M., Raya, A., Kawakami, Y., Wang, J., Zhang, X., Burch, J., Kubalak, S. W., Kaliman, P., Belmonte, J. C. I., et al. (2005). Epicardial retinoid X receptor α is required for myocardial growth and coronary artery formation. Proceedings of the National Academy of Sciences of the United States of America 102, 18455–18460.

Perez-Pomares, J. M., Carmona, R., Gonzalez-Iriarte, M., Macias, D., Guadix, J. A. and Munoz-Chapuli, R. (2004). Contribution of mesothelium-derived cells to liver sinusoids in avian embryos. Dev Dyn 229, 465–474.

Picelli, S., Faridani, O. R., Björklund, Å. K., Winberg, G., Sagasser, S. and Sandberg, R. (2014). Full-length RNA-seq from single cells using Smart-seq2. Nature Protocols 9, 171.

Platzer, A. (2013). Visualization of SNPs with t-SNE. PloS one 8, e56883–e56883.

Qiu, X., Hill, A., Packer, J., Lin, D., Ma, Y.-A. and Trapnell, C. (2017a). Single-cell mRNA quantification and differential analysis with Census. Nature methods 14, 309–315.

Qiu, X., Mao, Q., Tang, Y., Wang, L., Chawla, R., Pliner, H. and Trapnell, C. (2017b). Reversed graph embedding resolves complex single-cell developmental trajectories. bioRxiv, 110668.

Ratnadiwakara, M., Archer, S. K., Dent, C. I., Ruiz De Los Mozos, I., Beilharz, T. H., Knaupp, A. S., Nefzger, C. M., Polo, J. M. and Anko, M.-L. (2018). SRSF3 promotes pluripotency through Nanog mRNA export and coordination of the pluripotency gene expression program. eLife 7, e37419.

Red-Horse, K., Ueno, H., Weissman, I. L. and Krasnow, M. A. (2010). Coronary arteries form by developmental reprogramming of venous cells. Nature 464, 549.

Ruiz-Villalba, A., Ziogas, A., Ehrbar, M. and Pérez-Pomares, J. M. (2013). Characterization of Epicardial-Derived Cardiac Interstitial Cells: Differentiation and Mobilization of Heart Fibroblast Progenitors. PLoS ONE 8, e53694.

Stuart, T., Butler, A., Hoffman, P., Hafemeister, C., Papalexi, E., Mauck, W. M., Stoeckius, M., Smibert, P. and Satija, R. (2018). Comprehensive integration of single cell data. bioRxiv, 460147.

Tandon, P., Miteva, Y. V., Kuchenbrod, L. M., Cristea, I. M. and Conlon, F. L. (2013). Tcf21 regulates the specification and maturation of proepicardial cells. Development (Cambridge, England) 140, 2409–2421.

Tian, X., Hu, T., Zhang, H., He, L., Huang, X., Liu, Q., Yu, W., He, L., Yang, Z., Yan, Y., et al. (2014). De novo formation of a distinct coronary vascular population in neonatal heart. Science 345, 90–94.

Tian, X., Hu, T., Zhang, H., He, L., Huang, X., Liu, Q., Yu, W., He, L., Yang, Z., Zhang, Z., et al. (2013). Subepicardial endothelial cells invade the embryonic ventricle wall to form coronary arteries. Cell Research 23, 1075.

Tirosh, I., Izar, B., Prakadan, S. M., Wadsworth, M. H., 2nd, Treacy, D., Trombetta, J. J., Rotem, A., Rodman, C., Lian, C., Murphy, G., et al. (2016). Dissecting the multicellular ecosystem of metastatic melanoma by single-cell RNA-seq. Science 352, 189–196.

Trapnell, C., Cacchiarelli, D., Grimsby, J., Pokharel, P., Li, S., Morse, M., Lennon, N. J., Livak, K. J., Mikkelsen, T. S. and Rinn, J. L. (2014). The dynamics and regulators of cell fate decisions are revealed by pseudotemporal ordering of single cells. Nat Biotechnol 32, 381–386.

Volz, K. S., Jacobs, A. H., Chen, H. I., Poduri, A., McKay, A. S., Riordan, D. P., Kofler, N., Kitajewski, J., Weissman, I. and Red-Horse, K. (2015). Pericytes are progenitors for coronary artery smooth muscle. eLife 4, e10036.

Wang, F., Flanagan, J., Su, N., Wang, L. C., Bui, S., Nielson, A., Wu, X., Vo, H. T., Ma, X. J. and Luo, Y. (2012). RNAscope: a novel in situ RNA analysis platform for formalin-fixed, paraffin-embedded tissues. J Mol Diagn 14, 22–29.

Wei, K., Díaz-Trelles, R., Liu, Q., Diez-Cuñado, M., Scimia, M.-C., Cai, W., Sawada, J., Komatsu, M., Boyle, J. J., Zhou, B., et al. (2015). Developmental origin of age-related coronary artery disease. Cardiovascular Research 107, 287–294.

Weinreb, C., Wolock, S., Tusi, B. K., Socolovsky, M. and Klein, A. M. (2018). Fundamental limits on dynamic inference from single-cell snapshots. Proc Natl Acad Sci U S A 115, E2467–E2476.

Xiao, Y., Hill, M. C., Zhang, M., Martin, T. J., Morikawa, Y., Wang, S., Moise, A. R., Wythe, J. D. and Martin, J. F. (2018). Hippo Signaling Plays an Essential Role in Cell State Transitions during Cardiac Fibroblast Development. Developmental Cell 45, 153–169.e156.

Zhang, H., Pu, W., Li, G., Huang, X., He, L., Tian, X., Liu, Q., Zhang, L., Wu, S. M., Sucov, H. M., et al. (2016). Endocardium Minimally Contributes to Coronary Endothelium in the Embryonic Ventricular Free Walls. Circ Res 118, 1880–1893.

Zhou, B., Ma, Q., Rajagopal, S., Wu, S. M., Domian, I., Rivera-Feliciano, J., Jiang, D., von Gise, A., Ikeda, S., Chien, K. R., et al. (2008). Epicardial progenitors contribute to the cardiomyocyte lineage in the developing heart. Nature 454, 109–113.

Zhou, B. and Pu, W. T. (2012). Genetic Cre-loxP assessment of epicardial cell fate using Wt1-driven Cre alleles. Circulation research 111, e276–e280.

Ziegenhain, C., Vieth, B., Parekh, S., Reinius, B., Guillaumet-Adkins, A., Smets, M., Leonhardt, H., Heyn, H., Hellmann, I. and Enard, W. (2017). Comparative Analysis of Single-Cell RNA Sequencing Methods. Molecular Cell 65, 631–643.e634.

