## Supplementary Data for "Spatiotemporal analysis reveals significant overlap of key proepicardial markers in the developing murine heart"

### Supplementary Information

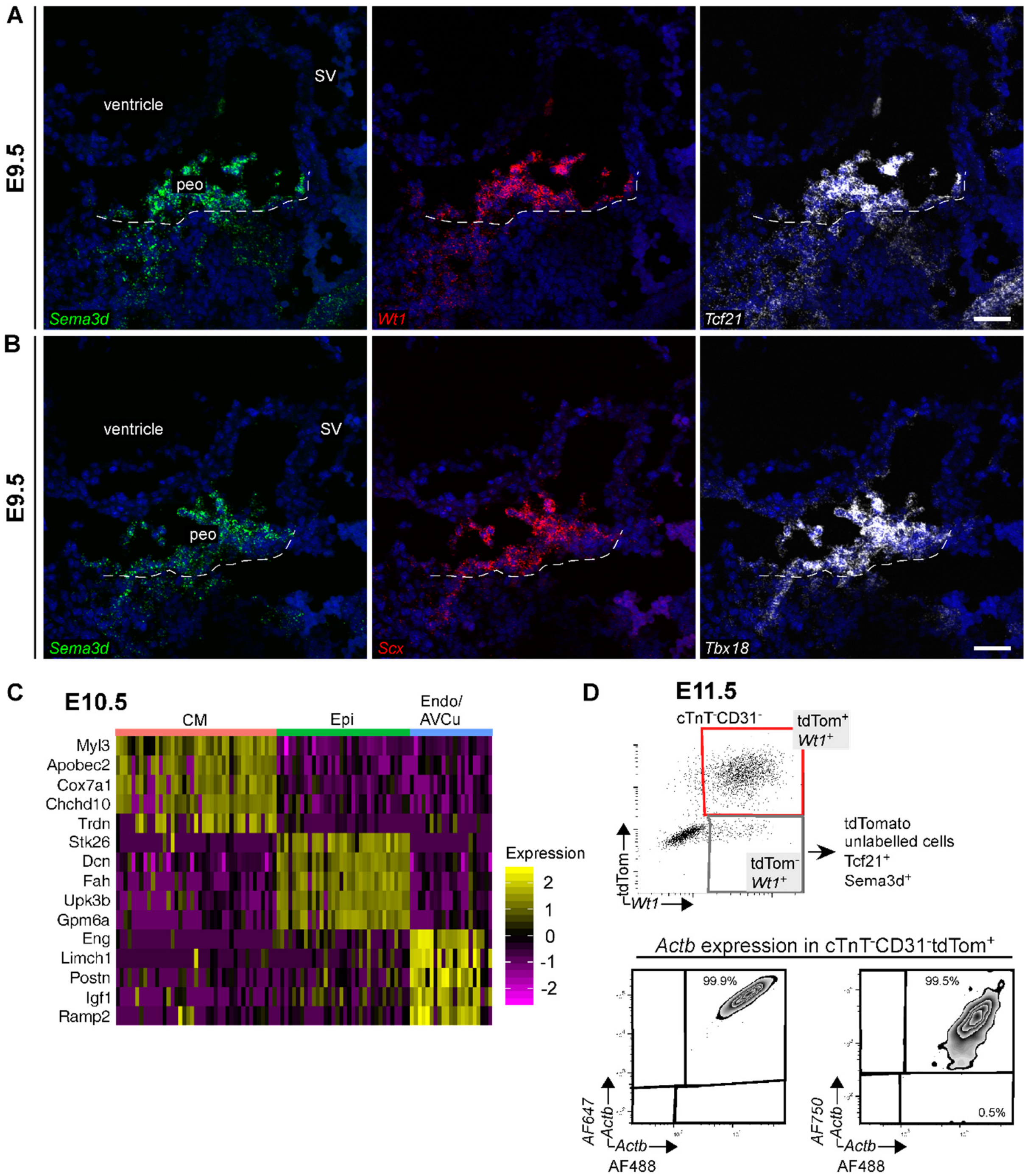

**Fig. S1. Gene expression of epicardial markers in the septum transversum region and confirmation of E10.5 scRNA-seq cluster identity.**

(A-B) RNA ISH staining of E9.5 embryo cryosections shows overlapping expression of *Sema3d*, (A) *Wt1*, *Tcf21*, (B) *Scx*, and *Tbx18* mRNA only in the proepicardial organ (dashed line) (n=5 embryos). (C) Heatmap of top 5 differentially expressed genes for each cluster in the E10.5 scRNA-Seq. High expression is indicated in yellow. (D) Flow cytometric analysis of *Wt1* and tdTomato at E11.5. Multiplex analysis of *Actb* expression in tdTomato+ cells. peo, proepicardial organ; SV, sinus venosus. Scale bars: 50µm in A-B.

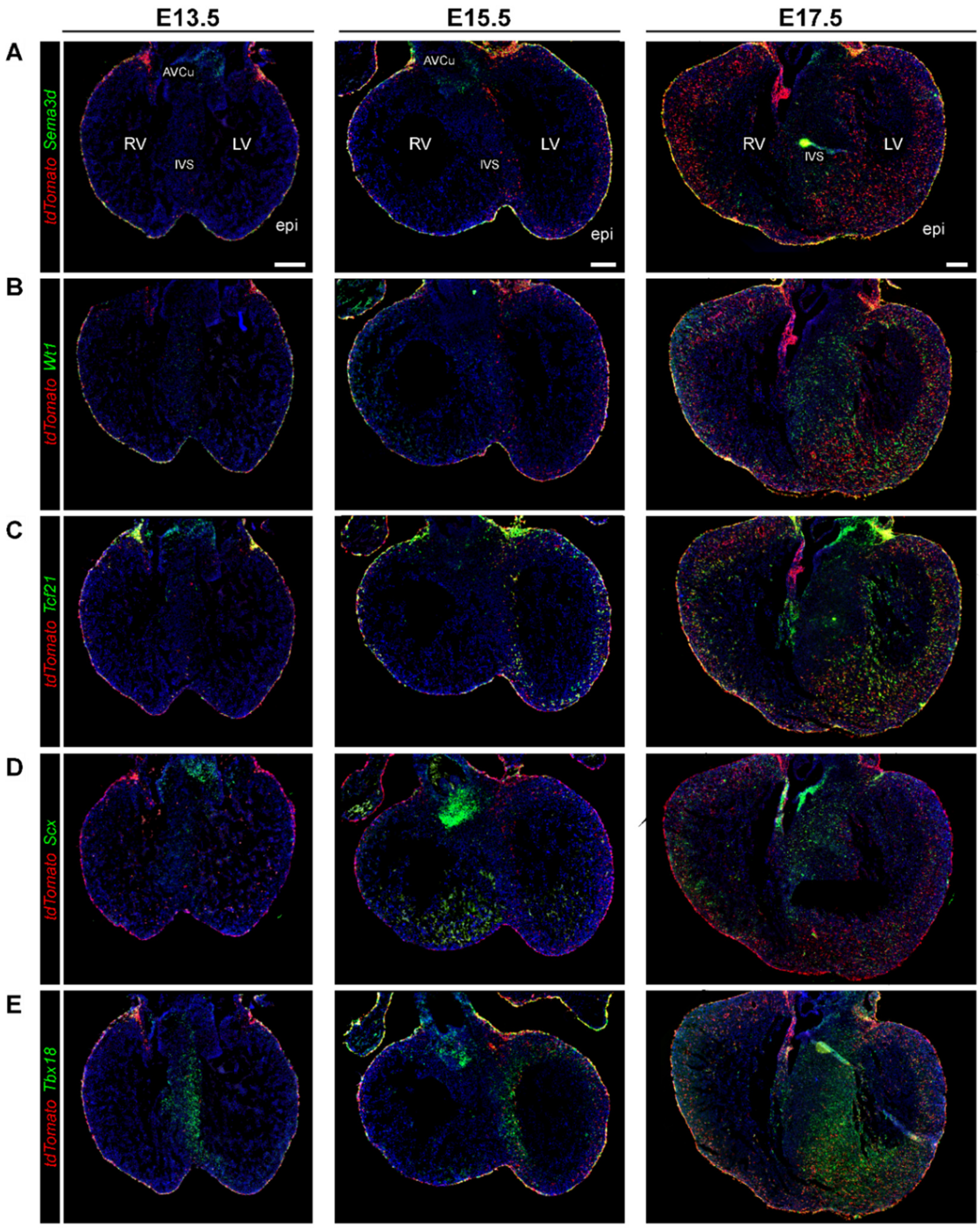

**Fig. S2. Non-specificity of the epicardial markers *Sema3d*, *Wt1*, *Tcf21*, *Scx*, and *Tbx18* at later stages of development.**

(A) RNA ISH co-staining of *Sema3d* (green) and *tdTomato* (red) at E13.5, E15.5, E17.5 shows overlap in the epicardium. *Sema3d* is also expressed in the atrioventricular cushion (AVCu) and in lymphatics. (B) RNA ISH co-staining of *Wt1* (green) and *tdTomato* (red) at E13.5, E15.5, E17.5 shows overlap in the epicardium. *Wt1* is also expressed in coronary endothelial cells starting E13.5. (C) RNA ISH co-staining of *Tcf21* (green) and *tdTomato* (red) at E13.5, E15.5, E17.5 shows overlap in the epicardium and in EPDCs. *Tcf21* is also expressed in AVCu and in interstitial fibroblasts. (D) RNA ISH co-staining of *Scx* (green) and *tdTomato* (red) at E13.5, E15.5, and E17.5 shows very low expression of Scx in the epicardium. *Scx* is highly expressed in the atrioventricular cushion (AVCu). (E) RNA ISH co-staining of Tbx18 (green) and tdTomato (red) at E13.5, E15.5, E17.5 shows overlap in the epicardium. *Tbx18* is also expressed in cardiomyocytes in the septum and left ventricle and in vascular smooth muscle cells in the aorta. AVCu, atrioventricular cushion; RV, right ventricle; IVS, intraventricular septum; LV, left ventricle; epi, epicardium. Scale bars: 200µm in A.

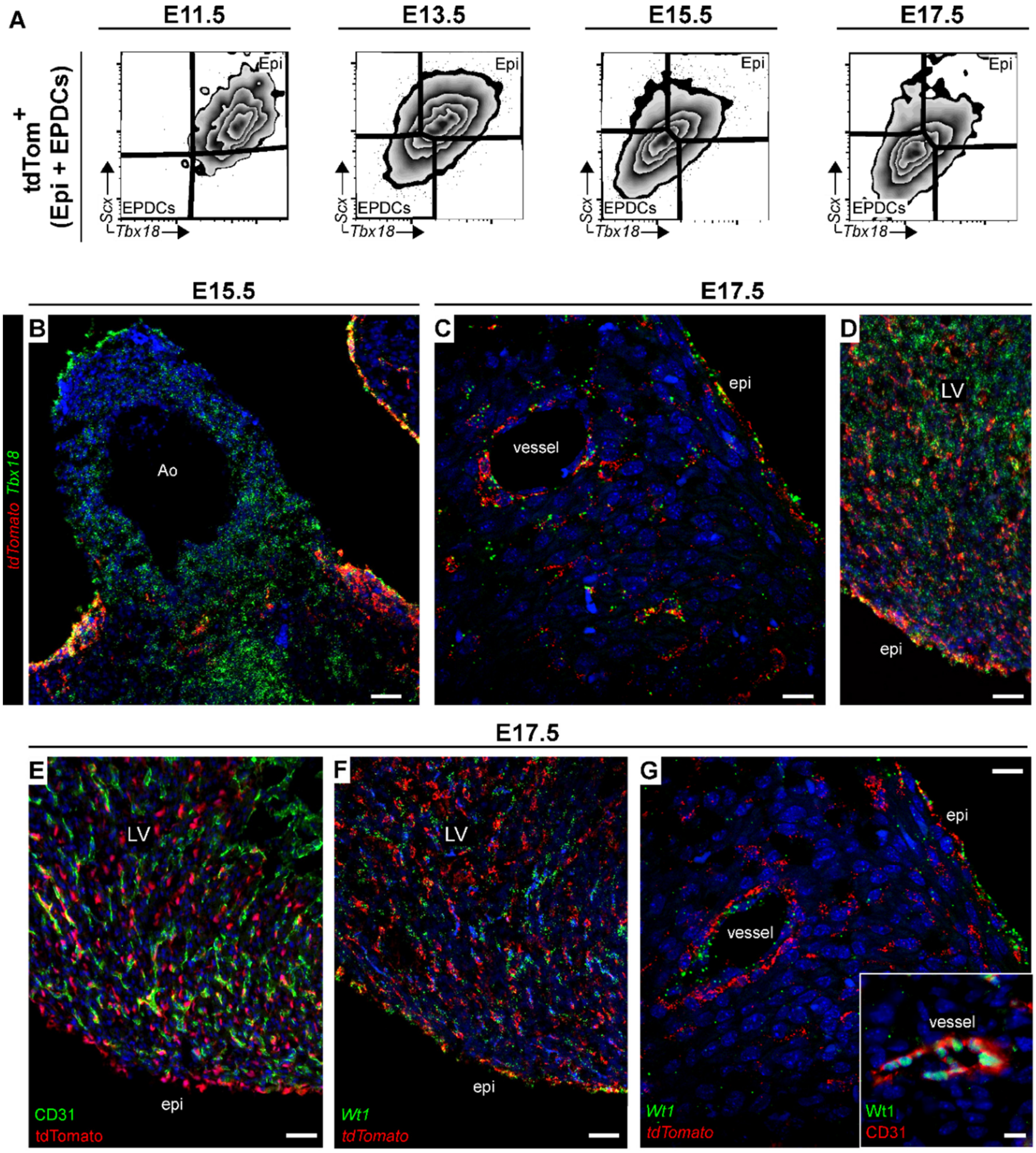

**Fig. S3. *Scx* and *Tbx18* expression decreases as epicardial cells transition to EPDCs and *Tbx18* and *Wt1* are expressed *de novo* in the coronary vasculature.**

(A) Flow cytometric analysis of *Scx* and *Tbx18* expression in tdTomato$^{+}$ cells, representing the epicardial lineage, shows downregulation of these markers in EPDCs over development. (B-C) RNA ISH co-staining of *Tbx18* (green) and *tdTomato* (red) reveals *Tbx18* expression in vascular smooth muscle cells derived from both non-epicardial and epicardial sources. (D) *Tbx18* is expressed in cardiomyocytes. (E) Co-immunostaining of CD31 and tdTomato shows no contribution of the epicardial lineage to coronary endothelium. (F-G) RNA-ISH co-staining of *Wt1* (green) and *tdTomato* (red) reveals *Wt1* expression in non-epicardial derived coronary endothelium cells. Co-immunostaining of CD31 and WT1 shows overlap in vessels.Epi, epicardium; EPDCs, epicardial-derived cells; Ao, aorta; LV, left ventricle. Scale bars: 50µm in B-G; 10µm in inset G.

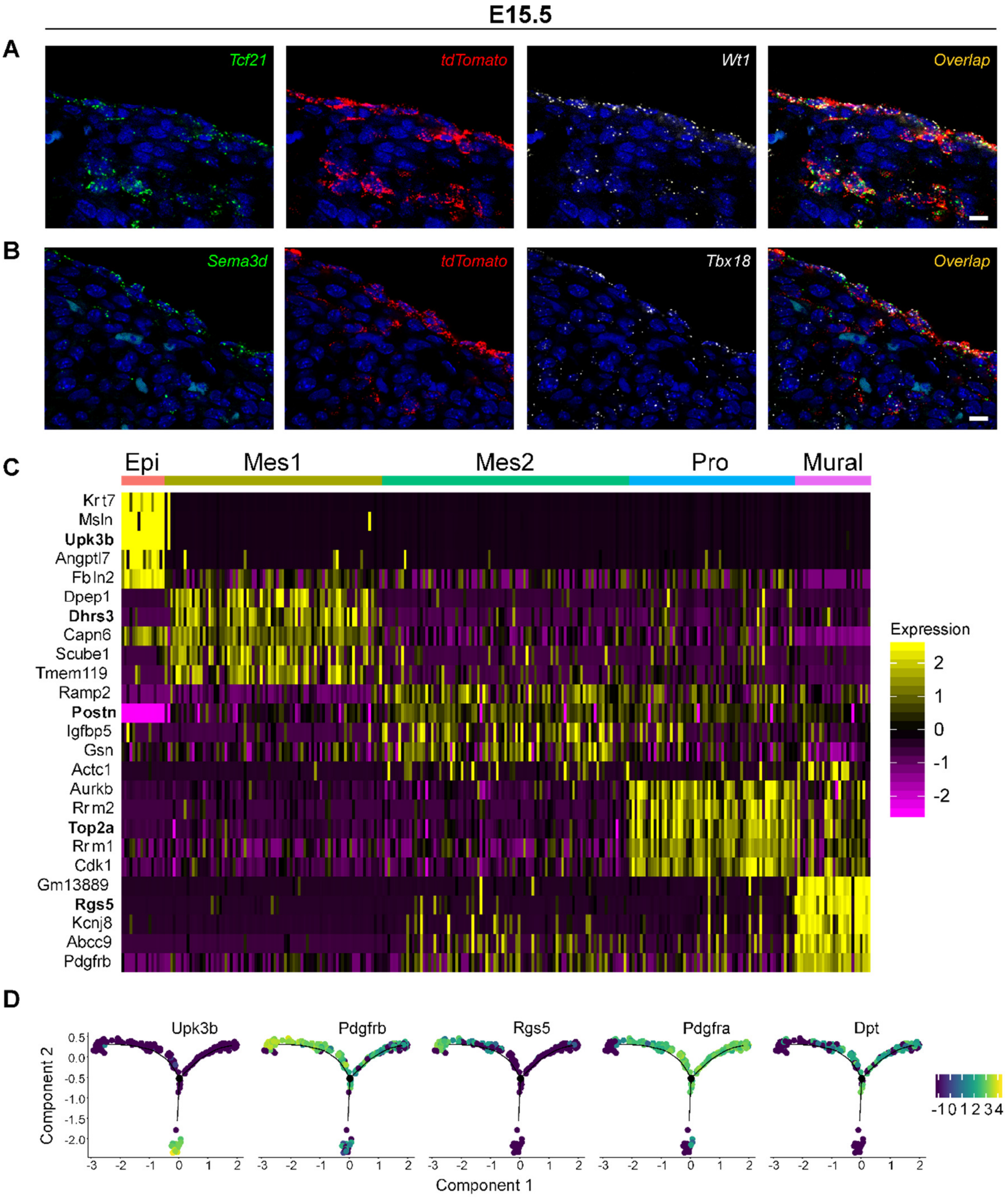

**Fig. S4. Expression of the selected epicardial markers at E15.5 and pseudotime representation of cell fate-associated genes.**

(A) RNA ISH co-staining of *Tcf21* (green), *tdTomato* (red) and *Wt1 (white)* reveals overlap in the epicardial lineage, albeit Tcf21 expression is low in the epicardium.(B) RNA ISH co-staining of *Sema3d* (green), *tdTomato* (red) and *Tbx18 (white)* reveals overlap in the epicardium and reduced expression in EPDCs.(C) Heatmap of top 5 differentially expressed genes per cluster in the E15.5 scRNA-Seq. High expression is indicated in yellow.(D) Visualisation of cluster marker genes in pseudotime reveals one epicardial origin ($Upk3b^{+}$) and two potential cell fates, one associated with mural genes (*Pdgfrβ* and *Rgs5*) and one associated with mesenchymal/fibroblast fate (*Pdgfrα* and *Dpt*).

**Table. S1: % Positive expressed (SED) in tdTom+ epicardial population at E11.5**

| | *Sema3d* | *Wt1* | *Tcf21* | *Tbx18* | *Scx* |
|---|---|---|---|---|---|
| Number of values | 3 | 3 | 3 | 3 | 3 |
| Minimum | 97.30 | 96.00 | 94.20 | 94.90 | 81.60 |
| 25% Percentile | 97.30 | 96.00 | 94.20 | 94.90 | 81.60 |
| Median | 97.70 | 98.20 | 94.90 | 98.60 | 90.10 |
| 75% Percentile | 98.90 | 98.20 | 97.40 | 99.50 | 94.40 |
| Maximum | 98.90 | 98.20 | 97.40 | 99.50 | 94.40 |
| **Mean** | **97.97** | **97.47** | **95.50** | **97.67** | **88.70** |
| Std. Deviation | 0.8327 | 1.270 | 1.682 | 2.438 | 6.514 |
| *Std. Error* | *0.4807* | *0.7333* | *0.9713* | *1.408* | *3.761* |

**Table. S2: % Positive expressed (SED) *Sema3d* in *Wt1⁺Tcf21⁺* epicardial population**

| | E11.5 | E13.5 | E15.5 | E17.5 |
|---|---|---|---|---|
| Number of values | 3 | 3 | 2 | 2 |
| Minimum | 98.20 | 99.50 | 99.20 | 96.20 |
| 25% Percentile | 98.20 | 99.50 | 99.20 | 96.20 |
| Median | 98.90 | 99.70 | 99.30 | 96.45 |
| 75% Percentile | 99.40 | 99.80 | 99.40 | 96.70 |
| Maximum | 99.40 | 99.80 | 99.40 | 96.70 |
| **Mean** | **98.83** | **99.67** | **99.30** | **96.45** |
| Std. Deviation | 0.6028 | 0.1528 | 0.1414 | 0.3536 |
| *Std. Error* | *0.3480* | *0.08819* | *0.1000* | *0.2500* |

**Table. S3: % Positive expressed (SED) *Scx* in *Tbx18⁺Sema3d⁺* epicardial population**

| | E11.5 | E13.5 | E15.5 | E17.5 |
|---|---|---|---|---|
| Number of values | 3 | 3 | 2 | 2 |
| Minimum | 82.80 | 88.70 | 97.10 | 93.20 |
| 25% Percentile | 82.80 | 88.70 | 97.10 | 93.20 |
| Median | 93.60 | 98.80 | 97.30 | 94.75 |
| 75% Percentile | 94.00 | 99.00 | 97.50 | 96.30 |
| Maximum | 94.00 | 99.00 | 97.50 | 96.30 |
| **Mean** | **90.13** | **95.50** | **97.30** | **94.75** |
| Std. Deviation | 6.354 | 5.890 | 0.2828 | 2.192 |
| *Std. Error* | *3.668* | *3.400* | *0.2000* | *1.550* |

**SED, Super-Enhanced Dmax Subtraction algorithm calculates percent positives when comparing histograms.
